# HIV-1-infected human macrophages, by secreting RANKL, contribute to enhanced osteoclastogenesis

**DOI:** 10.1101/2020.02.14.947614

**Authors:** Rémi Mascarau, Florent Bertrand, Arnaud Labrousse, Isabelle Gennero, Renaud Poincloux, Isabelle Maridonneau-Parini, Brigitte Raynaud-Messina, Christel Vérollet

## Abstract

HIV-1 infection is frequently associated with low bone density, which can progress to osteoporosis leading to a high risk of fractures. Only a few mechanisms have been proposed to explain the enhanced osteolysis in the context of HIV-1 infection. As macrophages are involved in bone homeostasis and are critical cell hosts for HIV-1, we asked whether HIV-1-infected macrophages could participate in bone degradation. Upon infection, human macrophages acquired some osteoclast features: they became multinucleated, upregulated the osteoclast markers RhoE and β3 integrin, and organized their podosomes as ring superstructures resembling osteoclast sealing zones. However, HIV-1 infected macrophages were not fully differentiated in osteoclasts as they did not upregulate NFATc-1 transcription factor and were unable to degrade bone. Investigating whether infected macrophages participate indirectly to virus-induced osteolysis, we showed that they produce RANKL, the key osteoclastogenic cytokine. RANK-L secreted by HIV-1-infected macrophages was not sufficient to stimulate multinucleation, but promoted the protease-dependent migration of osteoclast precursors. In conclusion, we propose that, by stimulating RANKL secretion, HIV-1-infected macrophages contribute to create a microenvironment that favors the recruitment of osteoclasts, participating to bone disorders observed in HIV-1^+^ patients.

## Introduction

Low bone density is frequent in HIV-1 infected patients and can progress to osteoporosis and high risk of fractures [1]. While lifespan of patients has significantly increased with antiretroviral therapy, longterm complications such as bone defects appeared. Although multiple factors, including antiretroviral therapy, contribute to bone loss in infected patients, bone deficits in non-treated patients attest for a role of the virus by-itself [2–5].

The skeleton is a dynamic organ undergoing continual remodeling thanks to the actions of bone-resorbing osteoclasts (OC), bone-forming osteoblasts and osteocytes. The balance between osteoblast and OC can be disturbed leading to bone defects. During HIV-1 infection, bone loss is associated with an increase in blood biomarkers for bone resorption, suggesting a major contribution of OC (*6,9*). While osteoblasts develop from cells of mesenchymal origin, OC originate from precursors derived from blood monocytes that fuse with embryonic osteoclasts and also from precursors that fuse together [6, 7]. OC are multinucleated giant cells that differentiate from fusion of myeloid precursors under the control of Macrophage Colony-Stimulating Factor (M-CSF) and Receptor Activator of Nuclear Factor-κB ligand (RANK-L) [6–8]. The degree of OC differentiation depends mainly on the activity of RANKL that is moderated by its physiological decoy receptor osteoprotegerin (OPG) [9]. OC differentiation mainly occurs through activation of the transcription factor Nuclear factor of activated T cells cytoplasmic 1 (NFATc1) [10]. In addition, proinflammatory cytokines such as tumor necrosis factor (TNF)-α, interleukin (IL)-6, IL-1β, and IL-17 favor tosteoclastogenesis, *i.e.* recruitment and/or differentiation of osteoclast precursors, and thus provide a supportive environment for osteoclastogenesis [11]. Terminally differentiated OC express high levels of the αvβ3 integrin adhesion receptor and of resorption-related enzymes, including cathepsin K (Ctsk), Tartrate Resistant Acidic Phosphatase (TRAP) and the subunit C1 of the V-type proton ATPase (ATP6v1c1). OC attachment to bone is mediated by an OC-specific structure called sealing zone. It is composed of a dense array of inter-connected F-actin structures, called podosomes that form a circular superstructure. Podosomes are present only in myeloid cells, including macrophages (MF), dendritic cells and OC [12]. The sealing zone anchors OC to the bone surface and creates a confined resorption environment, where protons and osteolytic enzymes are secreted [8, 10]. The formation of this structure is strongly regulated, involving the Src tyrosine kinase [13] and the small GTPase RhoE [14].

Today, only a few mechanisms have been proposed to explain the increase in osteolytic activity associated with HIV-1 infection comprising increased production of proinflammatory cytokines [15], disruption of the immune system [5, 16–18] and activation of the bone resorption activity of infected OC [19–21]. Through its action on T and B cells, HIV-1 infection leads to an increase in the RANKL/OPG ratio that stimulates OC differentiation [5, 17]. In addition, we have shown that HIV-1 infects osteoclasts precursors, enhancing their migration to bones and differentiation, and mature OC inducing modifications in the structure and function of the sealing zone. These changes markedly correlate with an enhanced OC adhesion and bone degradation capacity. This exacerbated osteolytic activity of infected OC is dependent on Src activation by the viral protein Nef [21].

Along with CD4 T lymphocytes, MF are critical cell hosts for HIV-1. Recent data highlight the capacity of MF to sustain active viral replication *in vivo,* their resistance to the viral cytopathic effects and their distribution in most tissues of the organism. MF can serve both as a vehicle for viral transmission, dissemination and cellular reservoir [22–27]. Moreover, MF play an important role in bone homeostasis and repair, involving a collaboration between infiltrating monocyte-derived MF and resident MF (also called osteal MF or osteomacs) [28]. Bone MF differ from OC in their specific expression of Siglec1 (CD169) [29], they are in close contact with osteoblasts, thus participating in the regulation of bone mineralization. They play critical role in immune responses to pathogens and to biomaterials, and also contribute to the induction, progression and resolution of fracture repair [28, 30, 31].

Thus, in order to further understand the mechanisms involved in the bone defects induced by HIV-1 infection, we challenge a novel hypothesis that proposes a role for HIV-1-infected MF in virus-induced osteolysis. We examined whether HIV-1 infection would induce a reprogrammation of MF towards an OC signature. Several arguments support this proposition: upon inflammatory conditions, myeloid cells such as dendritic cells can transdifferentiate into functional OC [32–34] and *in vitro*, HIV-1-infected MF acquire some OC characteristics (multinucleation, enhanced capacity to degrade organic matrices and, when seeded on glass, organization of their podosomes into circular structures) [26, 35].Another possibility would be an indirect effect of MF infection *via* a modification in secreted molecules, which may exert bystander effects on the recruitment or on differentiation of surrounding OC precursors. Here, we obtained evidence on the critical role of HIV-infected MF on the secretion of RANKL which supports osteoclastogenesis.

## Results

### HIV-1 infection of macrophages induce the expression of a subset of osteoclast markers

To determine whether HIV-1-infected MF acquired OC characteristics, human MF derived from primary monocytes were infected with two R5 MF-tropic strains, ADA and NLAD8. Ten days post-infection, both strains induced an efficient and productive infection as judged by the percentage of infected cells determined by immunofluorescence (IF) analysis with an antibody against the viral protein gag, by quantification of the intracellular mRNA level and the concentration of gag in cell supernatants (Supplemental Fig. 1A and Fig. 1A). As expected from previous data [35, 36], HIV-1 infection of MF with either ADA or NLAD8 strains triggered efficient MF fusion into multinucleated giant cells (MGC) compared to non-infected cells. Interestingly, in our experimental conditions, the fusion index of HIV-1-infected MF was similar to the one of OC (monocytes differentiated with M-CSF and RANKL from the same donor) (Fig. 1A-B). MGC formation was also observed after infection of MF with two clinical viral strains [37] (Supplemental Fig. 1B), even if the infectivity rate was lower (data not shown). Then, we analyzed the expression of genes which are specifically overexpressed during OC differentiation (mRNA level quantification normalized to non-infected MF differentiated from the same donor). We first verified that the genes encoding for the transcription factor *NFATc1*, the osteolytic enzymes *TRAP*, *Ctsk* and A*TP6v1c1* as well as the β3 integrin subunit (*ITGB3*) and *RhoE* were induced in OC compared to non-infected MF. In HIV-1-infected MF (ADA and NLAD8 strains) compared to uninfected MF, the level of *ITGB3* and *RhoE* expression was increased by 2- and 4-fold respectively, whereas the one of *Nfatc1*, *TRAP*, *Ctsk* and *ATP6v1c1* was not significantly modified (Fig. 1C). Western blot analyses showed that the variations in mRNA expression level translated into increased protein expression of β3 integrin and RhoE in MF infected with both viral strains compared to non-infected MF (Fig. 1D and data not shown). Consistently with RT-qPCR analysis, no change was observed in the expression level of Ctsk protein (Supplemental Fig. 1C).

**Figure 1:**
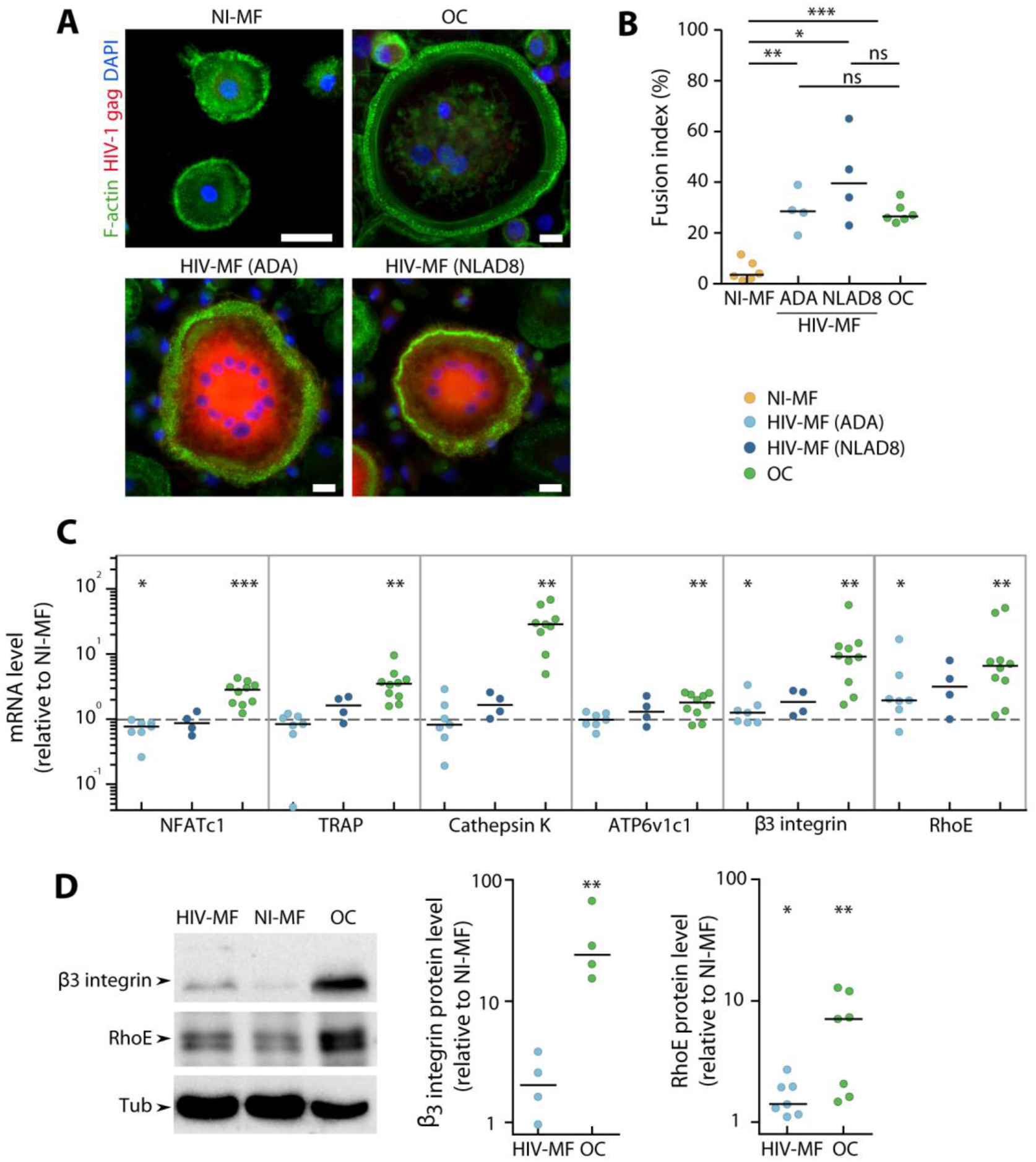
HIV-1 infection induces macrophage (MF) fusion and the expression of some osteoclast (OC) markers. MF were infected or not with HIV-1 (indicated strains), and compared to autologous OC for cell fusion (A-B) and expression of different OC markers (C-D). (A) Representative immunofluorescence (IF) images of uninfected MF (NI-MF), MF infected with HIV-1 (ADA or NLAD8 strain), and OC after staining for HIV-gag (red), F-actin (green), and nuclei (DAPI, blue). Scale bar, 10 μm. (B) Quantification of the fusion index evaluated by IF, corresponding to the percentage of nuclei within multinucleated cells. Bars represent median, n = 4 to 6 donors, 300 cells analyzed per donor and per condition. (C) mRNA expression of genes overexpressed in OC assayed by RT-PCR using the ΔΔCT method in MF infected with HIV-1 and in OC. Actin mRNA level was used as control. Values are normalized to mRNA level in autologous uninfected MF. Bars represent median, n = 4 to 10 donors. (D) Non infected MF (NI-MF), MF infected with HIV-1 NLAD8 strain and OC lysates were subjected to Western blot using antibodies against β3-integrin, RhoE and α-Tubulin as loading control. A representative blot (left panel) and quantification of the protein level ratio over autologous NI-MF (right panel) are shown. Bars represent median, n = 4 to 7 donors. * p ≤0.05; ** p ≤0.01; *** p ≤0.001, ns: not significantly different.

Together, these results indicate that HIV-1 infection of MF induces multinucleation and the expression of OC cytoskeletal markers, *i.e*. RhoE and β3 integrin, but is not sufficient to trigger the expression of all characteristic genes/proteins of OC.

### HIV-1 infection of macrophages induces the formation of sealing zone-like structures

As HIV-1 infected MF showed a partial differentiation toward OC, we next characterized the consequences of infection on the organization of podosomes, the main F-actin structure in MF and OC [12]. When cells were seeded on glass, most uninfected MF exhibited scattered podosomes while the majority of HIV-1(NLAD8)-infected MF reorganized their podosomes into clusters, rings or belts, as described for OC at different stages of differentiation [38] (Fig. 2A). Similar results were obtained upon infection of MF with the ADA strain [35]. The capacity of HIV-1-infected MF to form actin circular structures on glass was, at least, equivalent to the one of OC from the same donor. When cells were seeded on bone slices, we observed that about 15% of HIV-1 infected MGC formed podosome superstructures whose size was systematically smaller than the one of OC sealing zone (Fig. 2B-C). Noninfected MF did not present any specific actin organization while 80% of OC formed a classical sealing zone (Fig. 2B-C and Supplemental Figure 2). When looking at vinculin, a protein accumulated in podosomes, we showed that this protein colocalized to the actin ring structure assembled in HIV-1-infected MF, as observed in OC sealing zone (Fig. 2D).

**Figure 2:**
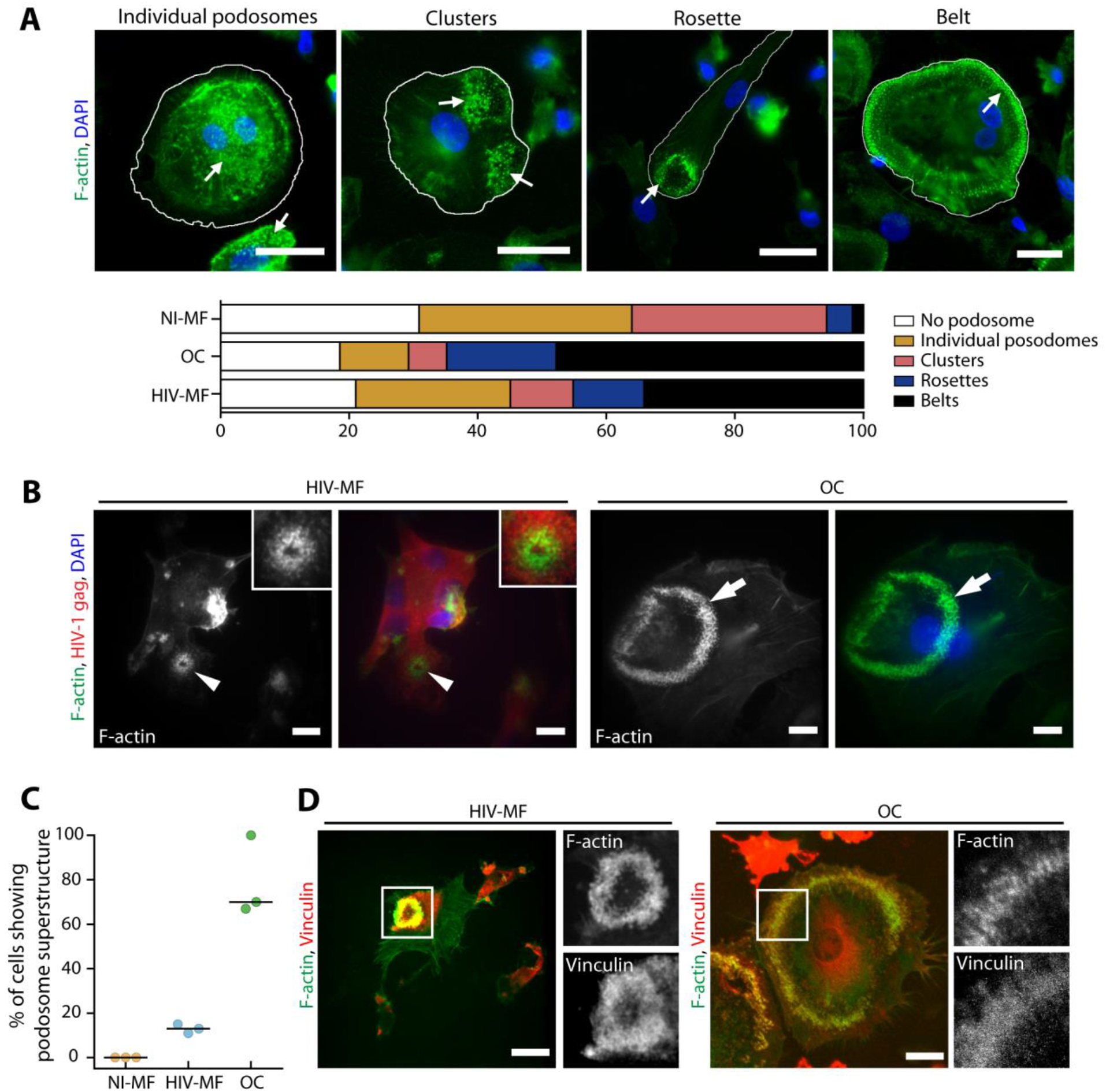
HIV-1 infection of MF promotes podosome organization into super-structures. (A) Top: Representative IF images of different podosome organizations in MF or OC seeded on glass coverslips, after staining for F-actin (green) and nuclei (DAPI, blue). White arrows show podosome structures. Scale bar, 20 μm. Bottom: Quantification of the different podosome organizations in MF, OC and HIV-1 infected-MF (HIV-MF, ADA strain), represented in percentage of total cells. n = 4 donors, 300 cells analyzed per donor. (B) Monocytes were seeded of bone slices and differentiated into MF or OC. Then, at day 7, MF were infected with HIV-1 (NLAD8 strain) and all cells were fixed at day 14. Left: representative IF images of podosome super structures in infected MF (arrowheads) and OC (arrows) stained for HIV-p24 (red), F-actin (green), and nuclei (DAPI, blue). Scale bar, 5 μm. Inserts show 2-fold magnification of the podosome structure in infected MF. (C) Quantification of the percentage of cells showing podosome organization on bones, evaluated by IF. Bars represent median, n = 3 donors, 100 cells analyzed per condition. (D) Representative image of podosome super-structures formed by infected-MF (HIV-MF, NLAD8 strain, left) or OC (right) seeded on bone slices, after staining for F-actin (green), vinculin (red) and nuclei (DAPI, blue). Scale bar, 10 μm. Inserts show magnification of white square.?

Thus, HIV-1 infected MF form F-actin structures sharing common characteristics with the OC sealing zone, even if number and size are lower.

### HIV-1 infection of macrophages does not enhance their bone degradation activity

As the formation of sealing zone is instrumental for osteolytic activity, we explored the capacity of HIV-1-infected MF to degrade the bone matrix. The bone resorption area observed for MF (close to 1% of the total bone surface) did not significantly increase upon HIV-infection (Fig. 3A-B), while it reached 15% for OC differentiated from the same donors. This parameter correlated with a low concentration of the C-terminal telopeptide of type 1 collagen (CTX) released in the supernatants of uninfected and infected MF, compared to the one obtained in OC supernatants which was around 50-fold higher (Fig. 3C). The morphology of resorption lacunae was examined by Scanning Electron Microscopy (SEM). As expected, OC formed long and large bone degradation trails [10, 21, 39]and Fig. 3D, upper panels). In the case of infected MF, we observed in some experiments (3 donors on 5) the rare appearance of individual round resorption pits referred to short trails; this degradation profile was never observed for uninfected MF. High magnification of the degraded pits showed superficial abrasion of bone microstructure in the case of infected MF compare to deep excavations with high porosity in the case of OC.

**Figure 3:**
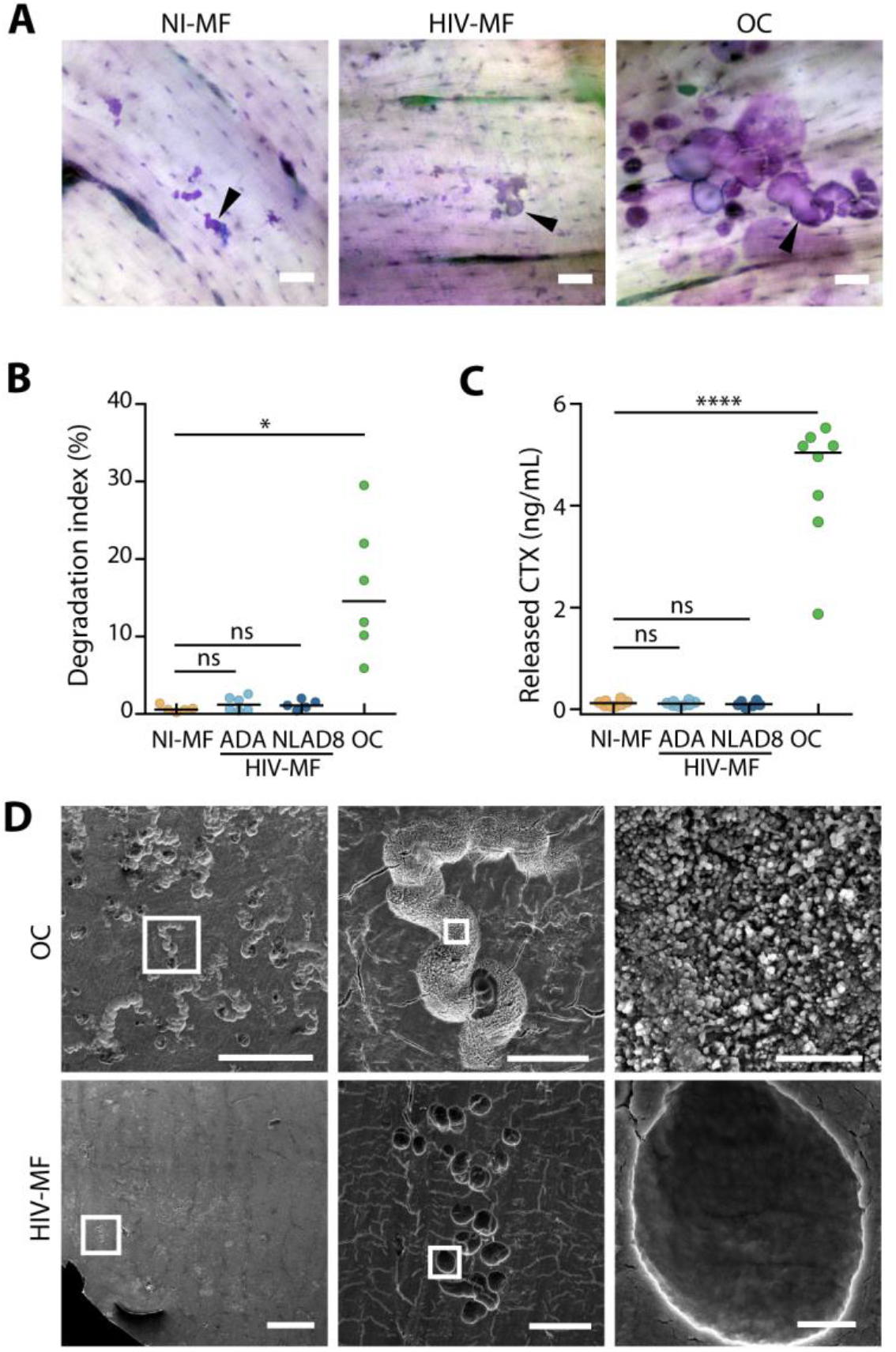
HIV-1 infection of MF does not enhance their bone degradation activity. Monocytes were seeded on bone slices and differentiated into MF or OC. At day 7, MF were infected with HIV-1 or not. At day 14, the supernatant was collected and cells were removed and bone slices were stained with Toluidine blue. (A) Representative images of the surface of bone slices cultured with OC and uninfected-(NI-MF) or infected-MF (HIV-MF, NLAD8 strain). Resorbed areas are revealed in purple (arrowheads). Scale bar, 10 μm. (B) Quantification of the resorbed area relative to total bone surface. Bars represent median, n = 5 to 6 donors. (C) Concentration of the bone degradation marker CTx in the supernatants was measured by ELISA. Bars represent median, n = 6 to 8 donors. (D) Representative scanning electron microscopy images showing bone resorption pits formed by OC or infected-MF (HIV-MF, NLAD8 strain). Scale bars indicated. * p ≤ 0.05; **** p ≤ 0.0001, ns: not significantly different.

All these data show that while HIV-1-infected MF acquire some OC characteristics such as multinuclearity and organized podosome rings on bones, they are not fully competent for bone resorption.?

### RANKL is secreted by infected macrophages and promotes 3D migration of osteoclast precursors

We then investigated whether HIV-infected MF through a modification in secreted molecules may exert bystander effects on the recruitment or differentiation of surrounding OC precursors. Indeed, an abundant literature reports the importance of cytokines in osteoclastogenesis, as they control both chemotaxis of OC precursors and OC differentiation *per se* [11]. First, the expression of the key osteoclastogenic cytokine, RANKL was examined. Interestingly, in conditioned medium of HIV-1-infected MF (CmHIV), RANKL concentration was increased in comparison with conditioned medium of uninfected control MF (CmCTL) (300 pg/mL of RANKL after infection *versus* 70 pg/mL) (Fig. 4A). The level of proinflammatory cytokines such as IL-1β, IL-6 and TNF-α, known to synergize with RANKL, were assessed [11]. The expression level of IL-1β and IL-6 remained below the threshold of detection in MF supernatants even after HIV-1 infection, and the level of TNF-α mRNA was not modified upon infection (Supplemental Fig. 3). Second, we investigated whether HIV-infected MF, by a bystander effect, could promote OC differentiation and bone degradation. To this end, OC precursors were differentiated in the presence of CmHIV or CmCTL, and the fusion index (as a parameter for OC differentiation) and bone degradation were measured. As infection of OC precursors promotes their differentiation [21], OC precursors were pre-treated, before addition of conditioned media, with maraviroc to prevent HIV-1 entry. The fusion of OC precursors and bone degradation were not significantly increased by incubation with CmHIV compared to CmCTL (Fig. 4B and data not shown). Finally, we tested whether CmHIV could impact the migration of OC precursors. The recruitment of OC precursors from blood to bones requires proteases *in vivo* [40], and we showed previously that defects in the 3 dimensional (3D) protease-dependent mesenchymal migration of these cells *in vitro* correlates with lower recruitment of OC to bones *in vivo* [21, 41]. In Matrigel, human OC precursors use the mesenchymal migration [42, 43]. We found that, using CmHIV as chemoattractant in the lower chamber, the percentage of cells infiltrating Matrigel was enhanced (Fig. 4C). This is not due to a potential infection of the cells inside the matrix by viral particles contained in CmHIV as addition of viral stock in the lower chamber has no effect on OC precursor migration. Importantly, as shown in Figure 4D, we also found an increase of the percentage of migrating OC precursors using as chemoattractant a range of recombinant RANKL concentrations close to the ones measured in CmHIV.

**Figure 4:**
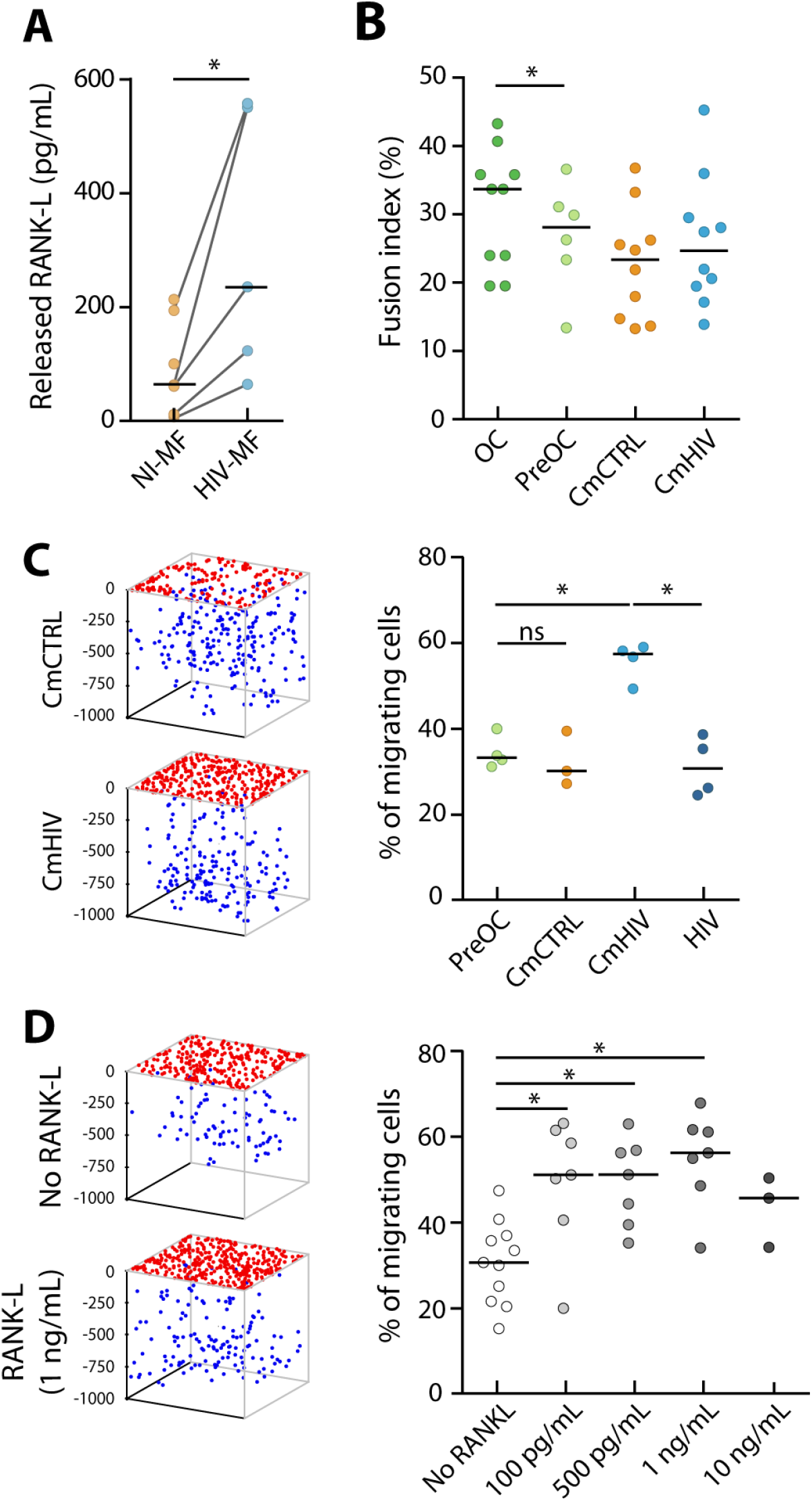
RANKL is secreted by HIV-1 infected MF and promotes 3D migration of OC precursors. (A) MF were infected with HIV-1 (HIV-MF, NLAD8 strain) for 7 days, and the level of released RANKL was measured by ELISA in the supernatant and compared to uninfected MF (NI-MF) from the same donor. Bars represent median, n = 5 to 8 donors. (B) Monocytes were differentiated for 3 days in presence of sub-optimal concentrations of RANKL and then exposed to the supernatant of infected (CmHIV) or uninfected (CmCTL) MF for additional 10 days. Cells were then fixed and the fusion index was quantified by IF. Bars represent median, n = 6 to 10 donors. (C-D) Supernatant of infected (CmHIV) or uninfected MF (CmCTL) treated or viral stock (supernatant of HEK293T transected with a pro-viral plasmid) (C), or different concentrations of RANKL (D) was added at the bottom of 3D Matrigel matrices. OC precursors were then seeded at the top of the matrices and allowed to migrate for 72h. Left: 3D representation of the positions of OC precursors that have migrated (blue dots) or not (red dots) in a representative migration assay. Right: quantification of the percentage of cells inside the matrix. Bars represent median. (C) n = 3 to 5 donors performed in triplicates. * p ≤ 0.05, ns: not significantly different.

In conclusion, CmHIV is not sufficient to promote differentiation of OC precursors *in vitro*, but it promotes their 3D mesenchymal migration, in particular through RANKL secretion, suggesting that HIV-1-infected MF could exert a bystander effect to favor active recruitment of OC precursors to bones.

## Discussion

The mechanisms underlying bone defects and exacerbation of osteolysis resulting from HIV-1 infection need to be further investigated. It is established that different MF subsets are present in bones, they are essential for maintenance of the bone architecture notably during tissue regeneration and inflammatory responses [28, 30, 31].[44, 45] Here, we studied whether MF could participate into osteolysis associated with HIV infection. HIV-1-infected human MF acquired some OC features but failed to resorb bone. Importantly, we report that however infection of MF enhanced their RANKL secretion that promotes osteoclasteogenesis.

Upon HIV-1 infection with different MF-tropic viral strains, MF acquired OC characteristics, such as multinucleation and ability to organize podosomes into circular structures, when seeded on glass [35] or bone (this study). The effect on fusion was also observed using transmitted/founder viral strains, which are single variants responsible for the viral transmission in patients. In addition, an increase in RhoE and β3 integrin levels, two proteins essential for OC podosome patterning into a functional sealing zone [10, 14], was detected in infected MF. When plated on bone matrices, infected MF organized their actin cytoskeleton into circular and vinculin-positive superstructures reminiscent of sealing zone. These superstructures were not present in non-infected MF (Supplemental Fig. 2). The Factin structures formed in HIV-1-infected MF promoted strong adhesion to the substrates (data not shown) but produced only superficial and limited abrasion of bone substrates. These sealing zone-like structures are smaller in size, resembling the podosome rosettes induced by activated Src kinases in fibroblasts [46–48]. Thus, in HIV-1-infected MF, the induction of a circular podosome organization associated with an increase in RhoE and β3 integrin levels are not sufficient for generating functional sealing zone. It is likely that they are unable to acidify mineral, as we have previously shown that HIV-1 infection increases the capacity of MF to degrade collagen matrices in 2D and in 3D [26]. In agreement with the inability of these structures to degrade bone, we did not notice any changes in the mRNA level of NFATc-1, the OC-master transcription factor [49, 50] and several genes under its control such as TRAP and Ctsk. Mature dendritic cells are able, in some specific conditions, to trans-differentiate into bone resorbing multinucleated OC Rivollier, 2004 #2706;Wakkach, 2008 #2931}[34]. This is not the case for fully differentiated MF in the context of HIV-1 infection. However, knowing the plasticity of haematopoietic progenitors, it would be interesting to consider this possibility at earlier stages of MF differentiation.

As the major stimulus for osteoclastogenesis is RANKL [11], we then determined whether HIV-1-infected MF could be an active source of soluble RANKL and thus potentially exert a bystander effect on recruitment and/or fusion of OC precursors. In CmHIV (supernatant of infected MF), we detected a 3-fold increase of RANKL concentration compared to CmCTL (supernatant of non-infected MF). This concentration is equivalent to the one produced by activated T cells, which are known as efficient RANKL producers [51]. Although MF were not described to be a classical source of RANKL, *in situ* hybridization have previously reported RANKL expression in MF in inflammatory contexts such as periodontal diseases [52, 53]. Increase in RANKL secretion by HIV-1-infected MF could result from either an increased production or a cleavage of the membranous forms [54].Thus, we propose that, in addition to osteoblasts, osteocytes, T and B lymphocytes, infected MF can be a novel source of RANKL. Others studies have already proposed that HIV-1 alters the secretion of osteoclastogenic regulatory factors. In HIV-transgenic rats and in sera from HIV-1 infected individuals, a marked increase in RANKL production together with a reduction in OPG production by lymphocytes has been reported [18]. In both models, the increase of RANKL/OPG ratio was correlated with a marked reduction in bone mineral density [5, 17, 18]. In contrast to RANKL, the levels of the inflammatory cytokines (IL-1β, IL-6 and TNF-α), described as cooperative factors of RANKL-dependent osteoclastogenesis, remained undetectable in infected MF.

Although CmHIV notably contained RANKL, we showed *in vitro* that incubation of OC precursors with CmHIV could not efficiently promote their multinucleation. However, it stimulated their migration in 3D Matrigel matrices. It is likely that the average concentration of RANKL secreted in the CmHIV (300 pg/mL) was under the threshold for triggering multinucleation of osteoclasts precursors *in vitro*. Actually, it is much lower than the one used to differentiate monocytes in OC *in vitro* (about 2 logs above). However, it is reasonable to propose that the increase in RANKL secretion by infected MF could contribute to enhanced osteolysis *in vivo* by favouring the recruitment of OC precursors to bones. Actually, we showed that the mesenchymal 3D migration of OC precursors is increased using, as chemo-attractant, recombinant RANKL concentrations equivalent to infected MF conditioned medium (range 100-1000 pg/mL). In CmHIV, RANKL probably mainly contributes to enhanced migration of OC precursors, although it was not excluded that other cytokines could participate to this induction [55]. We have reported that a correlation exists between enhanced mesenchymal migration of OC precursors and the number of OC in bones [41]. Therefore, we expect that the enhancement of OC precursor migration by the microenvironment generated by HIV-1-infected MF could favor OC recruitment to bones, and thus participate in bone disorders observed in infected patients. In accordance with this proposal, HIV-1-transgenic rats have a reduced bone mass as a consequence of an increased number of OC [18]. Furthermore, transgenic mice expressing the viral protein Nef, a key actor in the bone resorption activity of HIV-1-infected OC, exhibited a reduced bone density and a marked increase in OC compared with controls [21]. In conclusion, in addition to the mechanisms described previously by us and others [5, 17, 21], this study identifies a novel mechanism (*i.e.* RANKL secretion by HIV-1-infected MF) which likely participates to the high osteolysis associated with HIV-1 infection.

In bone, both MF derived from infiltrating monocytes and resident MF participate to tissue homeostasis [45]. It is conceivable that, like OC and OC precursors [21], MF or their precursors could be infected in the bone environment and, by secreting RANKL, contribute to a local supportive environment for osteoclastogenesis and, possibly, to an increase of circulating RANKL. We propose that the virus disrupts the RANKL/OPG equilibrium by stimulating the production of RANKL by T and B lymphocytes [5, 17]and also by MF (this study). To conclude, using a RANKL-dependent bystander mechanism, HIV-1-infected MF participate to the recruitment and differentiation of OC precursors that participate to bone disorders encountered in HIV-1+ patients.

## Material and methods

### Preparation of monocyte-derived macrophages and monocyte-derived osteoclasts

Buffy coats were provided by Etablissement Français du Sang, Toulouse, France, under contract 21/PLER/TOU/IPBS01/2013-0042. According to articles L1243-4 and R1243-61 of the French Public Health Code, the contract was approved by the French Ministry of Science and Technology (agreement number AC 2009-921). Written informed consents were obtained from the donors before sample collection. Monocytes from healthy subjects were isolated from buffy coats, seeded on glass coverslips (Marienfeld) or bone slices (ImmunoDiagnostic Systems) and differentiated into macrophages and osteoclasts (OC) as previously described [21]. Briefly, to obtain macrophages, purified CD14^+^ monocytes were differentiated for 7 days in RPMI-1640 medium (GIBCO) supplemented with 10% Fetal Bovine Serum (FBS, Sigma-Aldrich), L-glutamin (10 mM, Gibco), Penicillin-streptomycin (1%, Gibco) and human recombinant M-CSF (20 ng/mL, PeproTech).For OC differentiation, monocytes were cultured in complete medium supplemented with M-CSF (50 ng/mL) and RANKL (30 ng/mL, Miltenyi Biotech). The medium was replaced every 3 days with medium containing M-CSF (25 ng/mL) and RANKL (100 ng/mL). Macrophages (MF) and osteoclasts (OC) from the same donor were used. For migration experiments, OC precursors at day 3 of differentiation were used as in [21]. For differentiation of OC precursors by conditioned medium, monocytes were pre-exposed to sub-concentration of RANKL (10 ng/ml) during three days, then were cultured in the presence of CmCTL (conditioned medium of uninfected control MF) or CmHIV (conditioned medium of HIV-1-infected MF) supplemented with FBS (10%) during additional nine days. To prevent infection of precursors, Maraviroc (5 μM, Sigma) was added to the culture medium 30 minutes before adding conditioned medium.

### Viruses, HIV-1 infection and preparation of conditioned media

Proviral infectious clones of the macrophage-tropic HIV-1 (ADA and NLAD8) were kindly provided by Serge Benichou (Institut Cochin, Paris, France) while Transmitted /Founder strains (SUMA, cat# 11748 and THRO, cat# 11745) were obtained through the NIH AIDS Reagent Program, Division of AIDS, NIAID, NIH from Dr. John Kappes and Dr. Christina Ochsenbauer. Virions were produced by transient transfection of 293T cells with proviral plasmids, as previously described [26]. HIV-1 p24 antigen concentration of viral stocks was assessed by a home-made ELISA (see below). HIV-1 infectious units were quantified, as reported [56] using TZM-bl cells (Cat#8129, NIH AIDS Reagent Program, Division of AIDS, NIAID, NIH from Dr. John C. Kappes, and Dr. Xiaoyun Wu). Macrophages and OC were infected at MOI 0.5 (corresponding to 0.7 ng of p24 for 2.10^6^ cells). HIV-1 infection and replication were assessed 10 days post-infection by measuring the expression of Gag gene by RT-qPCR, the infection index calculated after p24 immunostaining and the p24 level in cell supernatants by ELISA.

For preparation of conditioned media, MF were infected with HIV-1 (NLAD8 strain) at a MOI of 0.5. The conditioned control medium (CmCTL) was obtained from uninfected MF from the same donor. After washing at day 1 post infection, culture media were collected at least at day 10 post infection, centrifuged to eliminate cell debris and aliquots were stored at −80°C.

### RNA extraction and qRT-PCR

Total RNA was extracted using ready-to-use TRIzol Reagent (Ambion, Life Technologies) and purified with RNeasy Mini kit (Qiagen) following manufacturer’s instructions. Complementary DNA was reverse transcribed from 1 μg total RNA with Moloney murine leukemia virus reverse transcriptase (Sigma) using dNTP (Promega) and random hexamer oligonucleotides (ThermoFisher) for priming. qPCR was performed using SYBR green Supermix (OZYME) in an ABI7500 Prism SDS Real-Time PCR Detection System (Applied Biosystems). The mRNA content was normalized to β-actin mRNA and quantified using the 2-ΔΔCt method. Primers used for cDNA amplification were purchased from Sigma and are listed in Supplemental Table 1.

### ELISA

Cytokine quantification was performed in cell supernatants by sandwich ELISA using kits from ELISAGenie (RANKL), BD Bioscience (IL-6) or Invitrogen (IL-1β) according to manufacturer’s instructions. Antibodies used are described in Supplemental Table 2. HIV-1 p24 concentration of viral stocks and p24 released by infected MF was measured by a previously described home-made sandwich ELISA [56] with NIH reagents, NIH AIDS Reagent Program, Division of AIDS, NIAID.

### Immunofluorescence microscopy

Immunofluorescence (IF) experiments were performed as described (6, 16). Briefly, cells were fixed with PFA 3.7% (Sigma), Sucrose 30mM in PBS (Gibco), permeabilized with Triton X-100 0,3% (Sigma) for 10 minutes, and saturated with PBS BSA 1% (Euromedex) for 30 minutes. Cells were incubated with primary antibodies diluted in PBS BSA 1% for 1 hour, washed and then incubated with corresponding secondary antibodies (LifeTechnologies), AlexaFluor488-or TexasRed-labeled phalloidin (Invitrogen) and DAPI (Sigma) in PBS BSA 1% for 30 minutes. Coverslips were mounted on a glass slide using Fluorescence Mounting Medium (Dako). Slides were visualized with a Leica DM-RB fluorescence microscope or on a FV1000 confocal microscope (Olympus). Images were processed with ImageJ and Adobe Photoshop softwares. Primary antibodies used are described in Supplemental Table 2. The HIV infection index (total number of nuclei in HIV-stained cells divided by total number of nuclei x 100) and fusion index (total number of nuclei in multinucleated cells divided by total number of nuclei x 100) were quantified, as in [35]. The number of cells with podosomes, podosome ring structures and sealing zones was quantified after phalloidin and vinculin staining.

### Scanning electron microscopy

Scanning electron microscopy observations were performed as previously described [21]. Briefly, following complete cell removal by immersion in water and scraping, bone slices were dehydrated in a series of increasing ethanol. Critical point was dried using carbon dioxide in a Leica EMCPD300. After coating with gold, bone slices were examined with a FEI Quanta FEG250 scanning electron microscope.

### Immunoblot analyses

Total protein lysates were extracted as previously described [57]. Total proteins were separated through SDS-polyacrylamide gel electrophoresis, transferred and immunoblotted overnight at 4°C with indicated primary antibodies. Primary antibodies used are described in Supplemental Table 2. Secondary antibodies were the following: anti-rabbit and anti-mouse IgG, HRP-linked Antibody (Cell Signaling Technology). Proteins were visualized with Amersham ECL Prime Western Blotting Detection Reagent (GE Healthcare). Chemiluminescence was detected with ChemiDoc Touch Imaging System (Bio-Rad Laboratories). Quantification of immunoblot intensity was performed using Image Lab (BioRad). Quantifications were normalized to tubulin.

### 3D migration assays

3D migration assays of OC precursors in Matrigel (10–12 mg/mL, BD Biosciences) were performed as described [21]. Recombinant RANKL was used as chemoattractant in the lower chamber at the indicated concentrations. Supernatant of infected (CmHIV) or uninfected (CmCTL) macrophage was supplemented with 20% FBS, incubated or not with recombinant OPG (500 ng/mL, Miltenyi Biotech.) for 30 minutes and added in the lower chamber for 3D migration assay. OC precursors were detached and resuspended in RPMI supplemented with 0.5 % FBS. 5.10^4^ cells were added at the top of each well and let to migrate for 72h. Cells were then fixed in 3.7% PFA for 1h and pictures were taken automatically with a 10X objective at constant intervals (z=30μm) using the motorized stage of an inverted microscope (Leica DMIRB, Leica Microsystems). Cells were counted using ImageJ software.

### Bone resorption assays

To assess bone resorption activity, monocytes were seeded on bovine cortical bone slices (IDS Nordic Biosciences, Paris, France) and differentiated into MF or OC. Following complete cell removal by several washes with water, bone slices were stained with toluidin blue (Sigma) to detect resorption pits under a light microscope (Leica DMIRB, Leica Microsystems). Surface of bone degradation areas were quantified manually with ImageJ software. Cross-linked C-telopeptide collagen I (CTX) concentrations were measured using betaCrosslaps assay (Immunodiagnostic System laboratory) in the culture medium of OC grown on bone slices [21].

### Statistical analysis

Information on the statistical tests used, and the exact values of n (donors) can be found in the Figure Legends. All statistical analyses were performed using GraphPad Prism 6.0 (GraphPad Software Inc.). The statistical tests were chosen according to the following. Two-tailed paired or unpaired t-test was applied on data sets with a normal distribution (determined using Kolmogorov-Smirnov test), whereas two-tailed Mann-Whitney (unpaired test) or Wilcoxon matched-paired signed rank tests were used otherwise. p<0.05 was considered as the level of statistical significance (* p≤0.05; ** p≤0.01; *** p≤0.001; **** p≤0.0001).

## Acknowledgments

The authors are grateful to M. Dupont for the technical help, S. Benichou for providing HIV-1 strains and M. Ben-Neji for macrophage and osteoclast preparation. The authors also acknowledge the multi-pathogen BSL3 facility at IPBS, the IPBS-TRI imaging facility and Isabelle Fourquaux, CMEAB-TRI, for her help with scanning electron microscopy preparation. We also thank the AIDS Research and Reference Reagent Program, Division of AIDS, NIAID for providing several tools. This work was supported by the *Centre National de la Recherche Scientifique*, *Université Paul Sabatier*, the *Agence Nationale de la Recherche* (ANR16-CE13-0005-01), the *Agence Nationale de Recherche sur le Sida et les hépatites virales* (ANRS2014-CI-2, ANRS2014-049, ANRS2018-2), the ECOS-*Sud* program (A14S01), the *Fondation pour la Recherche Médicale* (FRM DEQ2016 0334894), the *Fondation Bettencourt-Schueller* and *Human Frontier Science Program* (RGP0035/2016). R.M. is supported by the *Fondation Toulouse Cancer Santé*.

## Author Contributions

Conceptualization, CV, BRM and IMP; Methodology, RM, FB, AL, IG, RP, BRM and CV; Investigation, RM, FB, AL, IG, RP, BRM and CV; Validation, RM, BRM and CV; Formal Analysis, RM, BRM and CV; Writing – Original Draft Preparation, RM, BRM and CV; Writing – Review & Editing, RM, RP, IMP, BRM and CV; Visualization, RM and RP; Supervision, BRM and CV; Project Administration, IMP, BRM and CV; Funding Acquisition, IMP, BRM and CV.

## References

1. Cotter, A.G., and Mallon, P.W. (2014). The effects of untreated and treated HIV infection on bone disease. Curr Opin HIV AIDS 9, 17–26.

2. Bruera, D., Luna, N., David, D.O., Bergoglio, L.M., and Zamudio, J. (2003). Decreased bone mineral density in HIV-infected patients is independent of antiretroviral therapy. Aids 17, 1917–1923.

3. Gibellini, D., Borderi, M., De Crignis, E., Cicola, R., Vescini, F., Caudarella, R., Chiodo, F., and Re, M.C. (2007). RANKL/OPG/TRAIL plasma levels and bone mass loss evaluation in antiretroviral naive HIV-1-positive men. J Med Virol 79, 1446–1454.

4. Grijsen, M.L., Vrouenraets, S.M., Steingrover, R., Lips, P., Reiss, P., Wit, F.W., and Prins, J.M. (2010). High prevalence of reduced bone mineral density in primary HIV-1-infected men. Aids 24, 2233–2238.

5. Titanji, K., Vunnava, A., Sheth, A.N., Delille, C., Lennox, J.L., Sanford, S.E., Foster, A., Knezevic, A., Easley, K.A., Weitzmann, M.N., et al. (2014). Dysregulated B cell expression of RANKL and OPG correlates with loss of bone mineral density in HIV infection. PLoS Pathog 10, e1004497.

6. Jacome-Galarza, C.E., Percin, G.I., Muller, J.T., Mass, E., Lazarov, T., Eitler, J., Rauner, M., Yadav, V.K., Crozet, L., Bohm, M., et al. (2019). Developmental origin, functional maintenance and genetic rescue of osteoclasts. Nature 568, 541–545.

7. Kotani, M., Kikuta, J., Klauschen, F., Chino, T., Kobayashi, Y., Yasuda, H., Tamai, K., Miyawaki, A., Kanagawa, O., Tomura, M., et al. (2013). Systemic circulation and bone recruitment of osteoclast precursors tracked by using fluorescent imaging techniques. J Immunol 190, 605–612.

8. Teitelbaum, S.L. (2011). The osteoclast and its unique cytoskeleton. Ann N Y Acad Sci 1240, 14–17.

9. Teitelbaum, S.L. (2000). Bone resorption by osteoclasts. Science 289, 1504–1508.

10. Georgess, D., Machuca-Gayet, I., Blangy, A., and Jurdic, P. (2014). Podosome organization drives osteoclast-mediated bone resorption. Cell Adh. Migr. 8, 191–204.

11. Tanaka, S. (2017). RANKL-Independent Osteoclastogenesis: A Long-Standing Controversy. J Bone Miner Res 32, 431–433.

12. Wiesner, C., Le-Cabec, V., El Azzouzi, K., Maridonneau-Parini, I., and Linder, S. (2014). Podosomes in space: Macrophage migration and matrix degradation in 2D and 3D settings. Cell Adh. Migr. 8, 179–191.

13. Luxenburg, C., Addadi, L., and Geiger, B. (2006). The molecular dynamics of osteoclast adhesions. Eur J Cell Biol 85, 203–211.

14. Georgess, D., Mazzorana, M., Terrado, J., Delprat, C., Chamot, C., Guasch, R.M., Perez-Roger, I., Jurdic, P., and Machuca-Gayet, I. (2014). Comparative transcriptomics reveals RhoE as a novel regulator of actin dynamics in bone-resorbing osteoclasts. Mol Biol Cell 25, 380–396.

15. de Menezes, E.G., Machado, A.A., Barbosa, F., Jr., de Paula, F.J., and Navarro, A.M. (2016). Bone metabolism dysfunction mediated by the increase of proinflammatory cytokines in chronic HIV infection. J Bone Miner Metab.

16. Ofotokun, I., McIntosh, E., and Weitzmann, M.N. (2012). HIV: inflammation and bone. Curr HIV/AIDS Rep 9, 16–25.

17. Titanji, K., Vunnava, A., Foster, A., Sheth, A.N., Lennox, J.L., Knezevic, A., Shenvi, N., Easley, K.A., Ofotokun, I., and Weitzmann, M.N. (2018). T-cell receptor activator of nuclear factor-kappaB ligand/osteoprotegerin imbalance is associated with HIV-induced bone loss in patients with higher CD4+ T-cell counts. AIDS 32, 885–894.

18. Vikulina, T., Fan, X., Yamaguchi, M., Roser-Page, S., Zayzafoon, M., Guidot, D.M., Ofotokun, I., and Weitzmann, M.N. (2010). Alterations in the immuno-skeletal interface drive bone destruction in HIV-1 transgenic rats. Proc Natl Acad Sci U S A 107, 13848–13853.

19. Gohda, J., and al., e. (2015). HIV-1 replicates in human osteoclasts and enhances their differentiation in vitro. Retrovirology 12, 12.

20. Ofotokun, I., Titanji, K., Vikulina, T., Roser-Page, S., Yamaguchi, M., Zayzafoon, M., Williams, I.R., and Weitzmann, M.N. (2015). Role of T-cell reconstitution in HIV-1 antiretroviral therapy-induced bone loss. Nat Commun 6, 8282.

21. Raynaud-Messina, B., Bracq, L., Dupont, M., Souriant, S., Usmani, S.M., Proag, A., Pingris, K., Soldan, V., Thibault, C., Capilla, F., et al. (2018). Bone degradation machinery of osteoclasts: An HIV-1 target that contributes to bone loss. Proc Natl Acad Sci U S A 115, E2556–E2565.

22. Boliar, S., Gludish, D.W., Jambo, K.C., Kamng’ona, R., Mvaya, L., Mwandumba, H.C., and Russell, D.G. (2019). Inhibition of the lncRNA SAF drives activation of apoptotic effector caspases in HIV-1-infected human macrophages. Proc Natl Acad Sci U S A 116, 7431–7438.

23. Clayton, K.L., Collins, D.R., Lengieza, J., Ghebremichael, M., Dotiwala, F., Lieberman, J., and Walker, B.D. (2018). Resistance of HIV-infected macrophages to CD8(+) T lymphocyte-mediated killing drives activation of the immune system. Nat Immunol 19, 475–486.

24. Ganor, Y., Real, F., Sennepin, A., Dutertre, C.A., Prevedel, L., Xu, L., Tudor, D., Charmeteau, B., Couedel-Courteille, A., Marion, S., et al. (2019). HIV-1 reservoirs in urethral macrophages of patients under suppressive antiretroviral therapy. Nat Microbiol 4, 633–644.

25. Rodrigues, V., Ruffin, N., San-Roman, M., and Benaroch, P. (2017). Myeloid Cell Interaction with HIV: A Complex Relationship. Front Immunol 8, 1698.

26. Verollet, C., Souriant, S., Bonnaud, E., Jolicoeur, P., Raynaud-Messina, B., Kinnaer, C., Fourquaux, I., Imle, A., Benichou, S., Fackler, O.T., et al. (2015). HIV-1 reprograms the migration of macrophages. Blood 125, 1611–1622.

27. Sattentau, Q.J., and Stevenson, M. (2016). Macrophages and HIV-1: An Unhealthy Constellation. Cell Host Microbe 19, 304–310.

28. Batoon, L., Millard, S.M., Raggatt, L.J., and Pettit, A.R. (2017). Osteomacs and Bone Regeneration. Curr Osteoporos Rep 15, 385–395.

29. Batoon, L., Millard, S.M., Wullschleger, M.E., Preda, C., Wu, A.C., Kaur, S., Tseng, H.W., Hume, D.A., Levesque, J.P., Raggatt, L.J., et al. (2019). CD169(+) macrophages are critical for osteoblast maintenance and promote intramembranous and endochondral ossification during bone repair. Biomaterials 196, 51–66.

30. Bozec, A., and Soulat, D. (2017). Latest perspectives on macrophages in bone homeostasis. Pflugers Arch 469, 517–525.

31. Culemann, S., Gruneboom, A., Nicolas-Avila, J.A., Weidner, D., Lammle, K.F., Rothe, T., Quintana, J.A., Kirchner, P., Krljanac, B., Eberhardt, M., et al. (2019). Locally renewing resident synovial macrophages provide a protective barrier for the joint. Nature 572, 670–675.

32. Rivollier, A., Mazzorana, M., Tebib, J., Piperno, M., Aitsiselmi, T., Rabourdin-Combe, C., Jurdic, P., and Servet-Delprat, C. (2004). Immature dendritic cell transdifferentiation into osteoclasts: a novel pathway sustained by the rheumatoid arthritis microenvironment. Blood 104, 4029–4037.

33. Wakkach, A., Mansour, A., Dacquin, R., Coste, E., Jurdic, P., Carle, G.F., and Blin-Wakkach, C. (2008). Bone marrow microenvironment controls the in vivo differentiation of murine dendritic cells into osteoclasts. Blood 112, 5074–5083.

34. Laperine, O., Blin-Wakkach, C., Guicheux, J., Beck-Cormier, S., and Lesclous, P. (2016). Dendritic-cell-derived osteoclasts: a new game changer in bone-resorption-associated diseases. Drug Discov Today 21, 1345–1354.

35. Verollet, C., Zhang, Y.M., Le Cabec, V., Mazzolini, J., Charriere, G., Labrousse, A., Bouchet, J., Medina, I., Biessen, E., Niedergang, F., et al. (2010). HIV-1 Nef triggers macrophage fusion in a p61Hck- and protease-dependent manner. J Immunol 184, 7030–7039.

36. Kadiu, I., and Gendelman, H.E. (2011). Human immunodeficiency virus type 1 endocytic trafficking through macrophage bridging conduits facilitates spread of infection. J Neuroimmune Pharmacol 6, 658–675.

37. Baxter, A.E., Russell, R.A., Duncan, C.J., Moore, M.D., Willberg, C.B., Pablos, J.L., Finzi, A., Kaufmann, D.E., Ochsenbauer, C., Kappes, J.C., et al. (2014). Macrophage Infection via Selective Capture of HIV-1-Infected CD4(+) T Cells. Cell Host Microbe 16, 711–721.

38. Jurdic, P., Saltel, F., Chabadel, A., and Destaing, O. (2006). Podosome and sealing zone: specificity of the osteoclast model. Eur. J. Cell Biol. 85, 195–202.

39. Soe, K., and Delaisse, J.M. (2017). Time-lapse reveals that osteoclasts can move across the bone surface while resorbing. J Cell Sci 130, 2026–2035.

40. Blavier, L., and Delaisse, J.M. (1995). Matrix metalloproteinases are obligatory for the migration of preosteoclasts to the developing marrow cavity of primitive long bones. J Cell Sci 108 (Pt 12), 3649–3659.

41. Verollet, C., Gallois, A., Dacquin, R., Lastrucci, C., Pandruvada, S.N., Ortega, N., Poincloux, R., Behar, A., Cougoule, C., Lowell, C., et al. (2013). Hck contributes to bone homeostasis by controlling the recruitment of osteoclast precursors. FASEB J 27, 3608–3618.

42. Gui, P., Ben-Neji, M., Belozertseva, E., Dalenc, F., Franchet, C., Gilhodes, J., Labrousse, A., Bellard, E., Golzio, M., Poincloux, R., et al. (2018). The Protease-Dependent Mesenchymal Migration of Tumor-Associated Macrophages as a Target in Cancer Immunotherapy. Cancer Immunol Res.

43. Van Goethem, E., Poincloux, R., Gauffre, F., Maridonneau-Parini, I., and Le Cabec, V. (2010). Matrix Architecture Dictates Three-Dimensional Migration Modes of Human Macrophages: Differential Involvement of Proteases and Podosome-Like Structures. J Immunol 184, 1049–1061.

44. Michalski, M.N., and McCauley, L.K. (2017). Macrophages and skeletal health. Pharmacol Ther 174, 43–54.

45. Pettit, A.R., Chang, M.K., Hume, D.A., and Raggatt, L.J. (2008). Osteal macrophages: a new twist on coupling during bone dynamics. Bone 43, 976–982.

46. Cougoule, C., Carreno, S., Castandet, J., Labrousse, A., Astarie-Dequeker, C., Poincloux, R., Le Cabec, V., and Maridonneau-Parini, I. (2005). Activation of the lysosome-associated p61Hck isoform triggers the biogenesis of podosomes. Traffic 6, 682–694.

47. Destaing, O., Sanjay, A., Itzstein, C., Horne, W.C., Toomre, D., De Camilli, P., and Baron, R. (2008). The tyrosine kinase activity of c-Src regulates actin dynamics and organization of podosomes in osteoclasts. Mol Biol Cell 19, 394–404.

48. Luxenburg, C., Parsons, J.T., Addadi, L., and Geiger, B. (2006). Involvement of the Src-cortactin pathway in podosome formation and turnover during polarization of cultured osteoclasts. J Cell Sci 119, 4878–4888.

49. Takayanagi, H. (2007). The role of NFAT in osteoclast formation. Ann N Y Acad Sci 1116, 227–237.

50. Takayanagi, H., Kim, S., Koga, T., Nishina, H., Isshiki, M., Yoshida, H., Saiura, A., Isobe, M., Yokochi, T., Inoue, J., et al. (2002). Induction and activation of the transcription factor NFATc1 (NFAT2) integrate RANKL signaling in terminal differentiation of osteoclasts. Dev Cell 3, 889–901.

51. Kawai, T., Matsuyama, T., Hosokawa, Y., Makihira, S., Seki, M., Karimbux, N.Y., Goncalves, R.B., Valverde, P., Dibart, S., Li, Y.P., et al. (2006). B and T lymphocytes are the primary sources of RANKL in the bone resorptive lesion of periodontal disease. Am J Pathol 169, 987–998.

52. Chen, B., Wu, W., Sun, W., Zhang, Q., Yan, F., and Xiao, Y. (2014). RANKL expression in periodontal disease: where does RANKL come from? Biomed Res Int 2014, 731039.

53. Liu, D., Xu, J.K., Figliomeni, L., Huang, L., Pavlos, N.J., Rogers, M., Tan, A., Price, P., and Zheng, M.H. (2003). Expression of RANKL and OPG mRNA in periodontal disease: possible involvement in bone destruction. Int J Mol Med 11, 17–21.

54. Hikita, A., and Tanaka, S. (2007). Ectodomain shedding of receptor activator of NF-kappaB ligand. Adv Exp Med Biol 602, 15–21.

55. Xu, Y., Kulkosky, J., Acheampong, E., Nunnari, G., Sullivan, J., and Pomerantz, R.J. (2004). HIV-1-mediated apoptosis of neuronal cells: Proximal molecular mechanisms of HIV-1-induced encephalopathy. Proc Natl Acad Sci U S A 101, 7070–7075.

56. Souriant, S., Balboa, L., Dupont, M., Pingris, K., Kviatcovsky, D., Cougoule, C., Lastrucci, C., Bah, A., Gasser, R., Poincloux, R., et al. (2019). Tuberculosis Exacerbates HIV-1 Infection through IL-10/STAT3-Dependent Tunneling Nanotube Formation in Macrophages. Cell Rep 26, 3586–3599 e3587.

57. Bouissou, A., Proag, A., Bourg, N., Pingris, K., Cabriel, C., Balor, S., Mangeat, T., Thibault, C., Vieu, C., Dupuis, G., et al. (2017). Podosome Force Generation Machinery: A Local Balance between Protrusion at the Core and Traction at the Ring. ACS Nano 11, 4028–4040.

