## Supplementary material for "HIV-1-infected human macrophages, by secreting RANKL, contribute to enhanced osteoclastogenesis": Supplental material

**Supplemental Tables**

**Supplemental table 1 :** list of primers used for cDNA amplification

| Gene | Forward primer (5'-3') | Reverse primer (5'-3') |
|---|---|---|
| Actin | TCCCTGGAGAAGAGCTACGA | AGGAAGGAAGGCTGCAAGAG |
| ATP6V1C1 | CGGCAACTTCAAAGAACAAT | AAGCCCAACAGGAACCACACTG |
| Cathepsin K | GATGACTGGACTCAAAGTACC | AAGCCCAACAGGAACCACACTG |
| Gag | AGTGGGGGGACATCAAGCAGCCATGCAAT | TGCTATGTCACTTTCCCCTTGGTTCTCT |
| NFATc1 | CACCGCATCACAGGGAAGAC | GCACAGTCAATGACGGCTC |
| RhoE | GACACTTCGGGTTCTCCT | CAAAGCAAATCAGCACAGC |
| TNFα | GAGGCCAAGCCCTGGTATG | CGGGCCGATTGATCTCAGC |
| TRAP | TGACTTCCTCAGCCAGCA | AGCCCACGCCATTCTCATCTTG |
| β3 integrin | CCTGCTCATCTGGAAACTC | TGGGTTGTTGGCTGTGTC |

**Supplemental table 2 :** list of primary antibodies used and applications

| Target protein | Host specie | Clonality | Supplier | Application | Catalog number |
|---|---|---|---|---|---|
| Anti-human HRP | Goat | Polyclonal | Sigma | ELISA | A0170 |
| HIV (detection) | Human | Polyclonal | NIH AIDS Reagent Program | ELISA | 3957 |
| HIV-p24 (Capture) | Mouse | Monoclonal (IgG1 κ), clone 183-H12-5C | NIH AIDS Reagent Program | ELISA | 3537 |
| Anti-mouse AF555 | Goat | Polyclonal | Cell Signaling | IF | 4084 |
| HIV-p24 | Mouse | Monoclonal (IgG1), clone FH190-1-1 | Beckman Coulter | IF | KC57-RD1 |
| Vinculin | Mouse | Monoclonal (IgG1), clone hVIN-1 | Sigma | IF | V9131 |
| Anti-mouse HRP | Goat | Polyclonal | Dako | WB | P0447 |
| Anti-rabbit HRP | Goat | Polyclonal | Dako | WB | P0448 |
| Cathepsin K | Rabbit | Polyclonal | Abcam | WB | ab19027 |
| RhoE | Mouse | Monoclonal (IgG1) | Cell Signaling | WB | 3664 |
| Tubulin | Mouse | Monoclonal (IgG1), clone B5-1-2 | Sigma | WB | T5168 |
| β3 integrin | Rabbit | Polyclonal | Cell Signaling | WB | 4702 |

**Supplemental Figure legends**

**Supplemental Figure 1 :** (A) Infection of MF with HIV-1 (ADA or NLAD8 strain) was evaluated by measuring the expression of the viral gene Gag by RT-qPCR (left panel, results are normalized to GAPDH expression), calculating the fusion index by immunofluorescence (middle) and measuring p24 release in the supernatant by ELISA (right). Bars represent median, n = 4 to 9 donors. (B ) Representative IF images of MF infected with Transmitted/founder strains SUMA (left) and THRO (right) after staining of HIV-p24 (red), F-actin (green), and nuclei (DAPI, blue). Scale bar, 20 µm. (E) Cathepsin K (Ctsk) protein expression level was measured by Western blot in lysates from OC and infected (HIV-MF , ADA or NLAD8 strain) or uninfected MF (NI-MF). Tubulin was used as loading control. A representative blot and quantification of CtsK level relative to autologous NI-MF are shown. Bars represent median, n = 6 donors. * $p \leq 0.05$ ; ** $p \leq 0.001$ ; *** $p \leq 0.001$, ns: not significantly different.

**Supplemental Figure 2 :** Monocytes were seeded on bone slices, differentiated into macrophages for 7 days. Cells were then infected or not with HIV-1 and all cells were fixed at day 14. Representative IF images of cells after staining for HIV-p24 (red), F-actin (green), and nuclei (DAPI, blue). Scale bar, 10 µm.

**Supplemental Figure 3 :** (A-B) MF were infected or not with HIV-1 (ADA or NLAD8 strain) for 10 days and their expression of pro-inflammatory cytokines was evaluated. (A) Released IL-1β (left) and IL6 (right) was measured in the supernatants by ELISA. Bars represent median, n = 5 to 7 donors. (B) TNFα expression was measured by RT-qPCR (results are normalized to actin). Bars represent median, n = 3 to 6 donors.
