## Supplementary figures and images for "HIV-1-infected human macrophages, by secreting RANKL, contribute to enhanced osteoclastogenesis"

### Supplental figures

##### Supplemental Figure 1

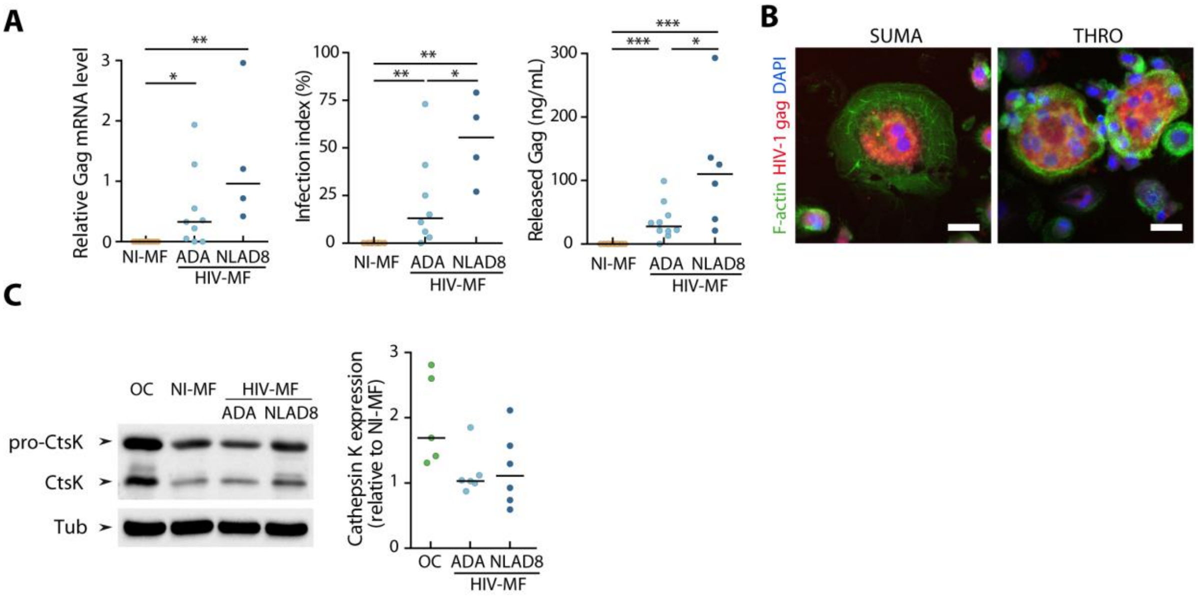

##### Supplemental Figure 2

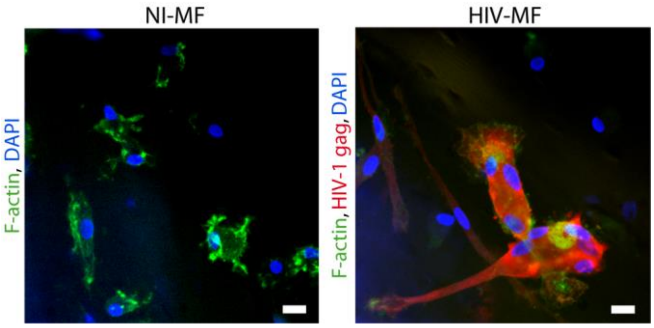

#### Supplemental Figure 3

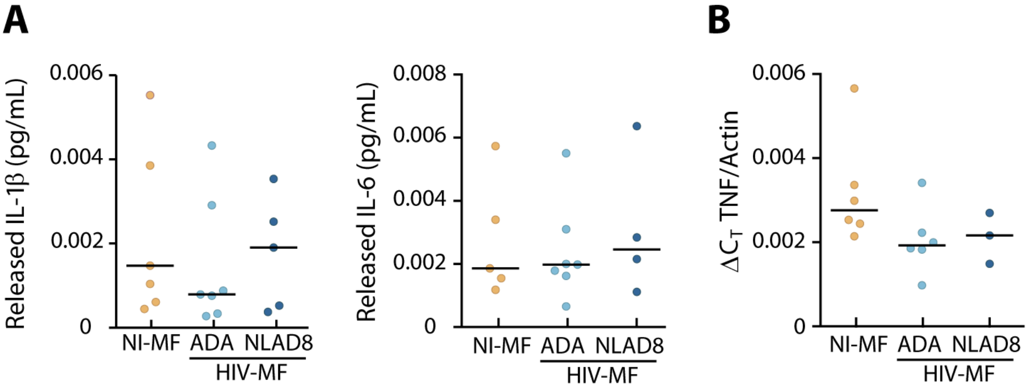
